# SARS-CoV-2 Envelope protein alters calcium signaling via SERCA interactions

**DOI:** 10.1101/2023.06.13.544745

**Authors:** Blanka Berta, Hedvig Tordai, Gergely L. Lukács, Béla Papp, Ágnes Enyedi, Rita Padányi, Tamás Hegedűs

## Abstract

The clinical management of severe COVID-19 cases is not yet well resolved. Therefore, it is important to identify and characterize cell signaling pathways involved in virus pathogenesis that can be targeted therapeutically. Envelope (E) protein is a structural protein of the virus, which is known to be highly expressed in the infected host cell and is a key virulence factor, however, its role is poorly characterized. The E protein is a single-pass transmembrane protein that can assemble into a pentamer forming a viroporin, perturbing Ca^2+^ homeostasis. Because it is structurally similar to regulins such as, for example, phospholamban, that regulate the sarco/endoplasmic reticulum calcium ATPases (SERCA), we investigated whether the SARS-CoV-2 E protein affects the SERCA system as an exoregulin. Using FRET experiments we demonstrate that E protein can form oligomers with regulins, and thus can alter the monomer/multimer regulin ratio and consequently influence their interactions with SERCAs. We also confirmed that a direct interaction between E protein and SERCA2b results in a decrease in SERCA-mediated ER Ca^2+^ reload. Structural modeling and molecular dynamics of the complexes indicates an overlapping interaction site for E protein and endogenous regulins. Our results reveal novel links in the host-virus interaction network that play an important role in viral pathogenesis and may provide a new therapeutic target for managing severe inflammatory responses induced by SARS-CoV-2.

## INTRODUCTION

The SARS CoV-2 coronavirus causes acute respiratory syndrome that emerged as a major global threat with high mortality (1). One of the four structural proteins, the envelope (E) protein plays an essential role in pathogenicity and virus replication (2). E protein is involved in viral assembly (3) and its deletion significantly decreases virulence (4). E protein has been found to be highly expressed in infected cells during the CoV replication cycle, also indicating that this protein is crucial for controlling cellular functions during replication (5). E protein is a small transmembrane (TM) protein of 75 amino acid (a.a.) residues with a single TM helix, terminated by a short N-terminal tail (1-8 a.a.) and a long C-terminal region (38-75 a.a.). The C-terminus of E protein harbors a PDZ binding motif (PBM) that interacts with human PDZ proteins, such as PALS1, perturbing cell polarity thus interfering with epithelial function (6). Removal of the PBM from SARS-CoV-1 E protein resulted in reduced expression of inflammatory cytokines and less pathogenicity in mice (7). SARS CoV-1 and CoV-2 E proteins differ only in three amino acids and one residue deletion. Since their C-terminal region is ordered (PDB IDs: 2mm4 and 5×29) (8,9) and disordered (10) in the presence and absence of lipids, respectively, it can be considered to contain a MemMoRF (Membrane Molecular Recognition Feature) (11).

Most of the E protein is distributed in the endoplasmic reticulum (ER) and the ER-Golgi intermediate compartment (ERGIC) (12). The E protein can form homopentamers acting as a viroporin that is permeable for cations including calcium, and was demonstrated to alkalinize ERGIC and the lysosome lumen (13).

The structure of E protein and its monomeric and pentameric forms closely resemble to the human regulins, which regulate the activity of the sarco/endoplasmic reticulum Ca^2+^ ATPase (SERCA) (14,15). The major function of these SERCA enzymes is to decrease the cytosolic calcium concentration, thus affecting calcium signaling by pumping calcium ions into the endoplasmic reticulum (ER) or sarcoplasmic reticulum (SR). SERCA proteins are encoded by three genes (ATP2A1, 2 and 3) that give rise to several isoforms, among which SERCA2b has the widest tissue expression pattern. This ubiquitous enzyme plays an essential housekeeping role in cellular calcium homeostasis, including that of lung parenchyma and vasculature. Therefore, SERCA has crucial functions in many cellular processes, such as proliferation, contractility, and apoptosis. The role of SERCA-dependent calcium transport in viral infections is receiving increasing attention (16–18). Novel data suggest that the vacuole membrane protein 1 (VMP1) interaction with viral proteins (19) results in inflammasome activation *via* SERCA pump activity modulation (20).

SERCA enzymes are modulated by binding of regulins such as phospholamban (PLN) or sarcolipin (SLN) (21). Some of these interactions reduce the Ca^2+^ affinity of SERCA pumps and the rate of Ca^2+^uptake into internal Ca^2+^stores. The two recently discovered regulins, endoregulin (ELN) and the another-regulin (ALN), inhibit the activity of SERCA as PLN (22,23). ALN exhibits a ubiquitous tissue expression pattern similar to SERCA2b (23), while ELN is expressed specifically in non-muscle tissues (epithelial cells of trachea and bronchi, lung and intestine) (23). Their SERCA inhibitory mechanisms are predicted to be similar to PLN due to their similar topology. PLN is a 52 amino acid (a.a.) long single-pass membrane protein containing three distinct parts. Its N-terminal cytoplasmic domain contains a MemMoRF (11) and this is followed by a TM helix and a short C-terminal tail located in the ER lumen. PLN was demonstrated to be in dynamic equilibrium between monomeric and pentameric states (24). The monomer is considered the “active” SERCA-inhibitory form while the pentameric form constitutes the “inactive” PLN pool (Figure 1). Oligomer formation was also observed for the other regulins, and these regulins were also bound to SERCA in a monomeric form (15). In addition, different regulins hetero-oligomerize with each other and this may lead to further fine-tuning of their effects on SERCA-dependent calcium transport (25).

**Figure 1.**
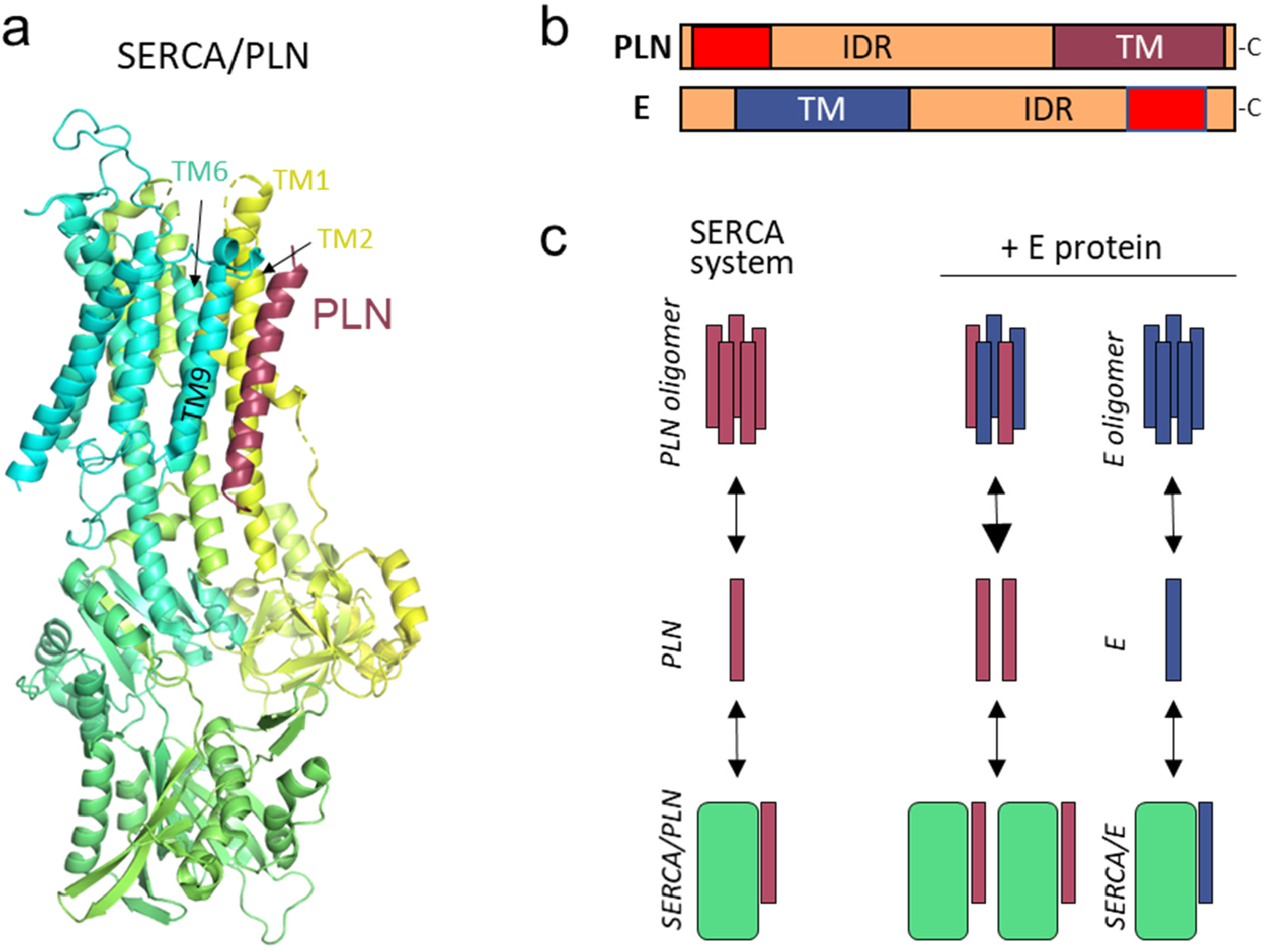
(**a**) The X-ray structure (PDBID: 4kyt) of the rabbit SERCA (yellow-green-turquoise) and PLN (burgundy) complex. (**b**) Topology of PLN and E proteins. TM: transmembrane helix; IDR: intrinsically disordered region; red: MemMoRF. (**c**) E protein may interfere with the SERCA system by perturbing the PLN pentamer pool or by interacting with SERCA. burgundy: PLN; blue: E protein; green: SERCA.

In this study, we investigated whether E protein can act as an exoregulin and interfere with the SERCA system, either by binding directly to SERCA or by interacting with endogenous regulins to modulate their monomer/multimer ratio. We also assessed if the interaction between E protein and SERCA2b has a functional relevance for Ca^2+^ signaling.

## RESULTS

### E protein co-localizes with SERCA2b and its regulins in the endoplasmic reticulum

To investigate the effect of E protein on the function of the SERCA-system, we coexpressed E protein with SERCA2b or regulins in HeLa cells. The well-known regulator, phospholamban and two non-muscle regulins, ALN and ELN were selected for our studies. The proteins were labeled with eGFP or mCherry. E protein was localized mainly in the endoplasmic reticulum and to a lesser extent in the ERGIC compartment at 24 h post-transfection (Supplementary Figure S1). SERCA was present in the ER that corresponds to its anticipated location (Figure 2). Although all regulins were detected in both the ER and the Golgi, their distribution between these two compartments was different. ALN exhibited distinct Golgi localization while PLN and ELN remained mainly in the ER even after 48 hours of transfection. E protein labeling indicated that it co-localized with all regulins and also with SERCA mostly in the ER (Figure 2).

**Figure 2.**
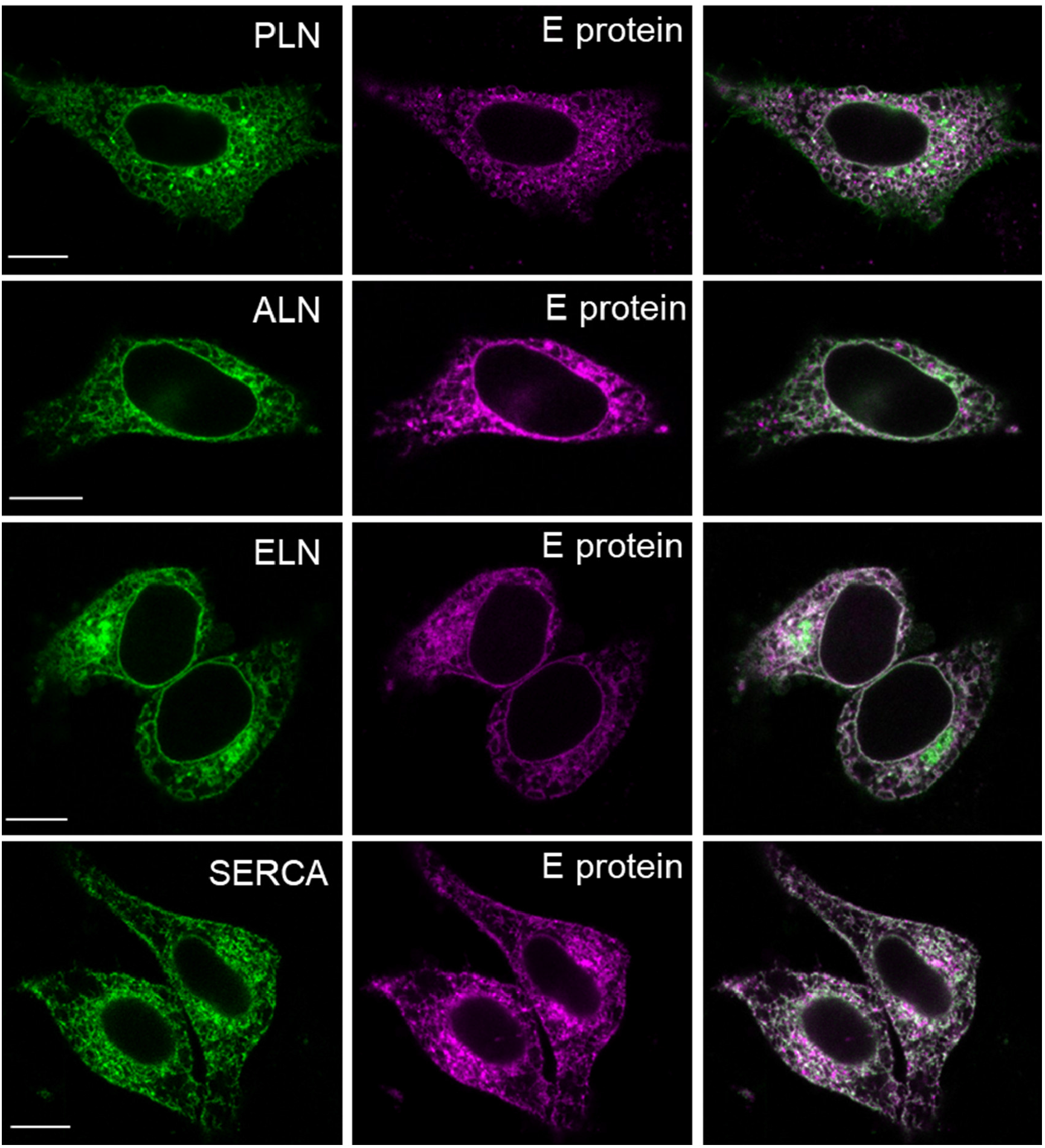
E protein colocalizes with SERCA and regulins in the endoplasmic reticulum. Columns with green, magenta, and white indicate the signals from localization of GPF and mCherry constructs and their colocalization, respectively. Scale bar is 10 μm.

### E protein forms heteromers with regulins and SERCA2b as detected by AP-FRET

Since co-localization does not necessarily indicate direct interaction, we used Förster Resonance Energy Transfer (FRET) to study the intermolecular association between the E protein and members of the SERCA/regulin system. FRET was monitored by acceptor photobleaching (AP-FRET) technique using donor eGFP and acceptor mCherry (Figure 3a). Homo-oligomerization of E protein and regulins is a well-known phenomenon, and we confirmed homo-oligomer formation of both E protein and regulins in our system (Figure 3b). The highest and lowest FRET values were observed for the PLN-PLN and ALN-ALN multimers, respectively. The lower ALN values, which still can be considered significant (26) when compared to control measurements with soluble eGFP and mCherry pairs, are probably due to the interfering effect of unlabeled endogenous ALN of HeLa cells.

**Figure 3.**
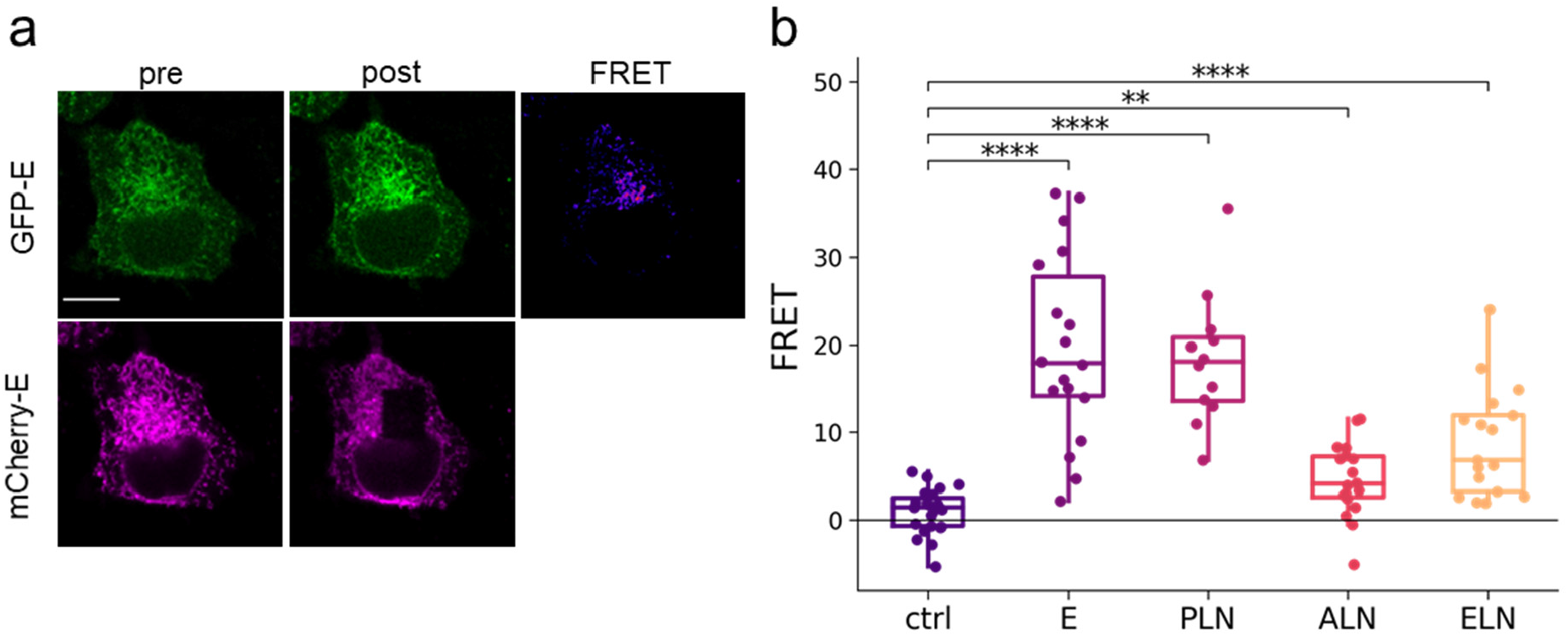
Homo-oligomerizations of E protein and regulins. (**a**) Fluorescence images of HeLa cells expressing GFP-(green) and mCherry-tagged (magenta) E protein. The first and second columns show images before (pre) and after (post) photobleaching, respectively. The FRET image shows FRET values calculated by *FRETcalc*. The scale bar is 10 μm. (**b**) FRET efficiency values produced by homo-oligomers. Median and interquartile ranges are indicated with box plots. Data were analyzed by Kolmogorov-Smirnov test (* p<0.05, ** p<0.01, *** p<0.001, ****p<0.0001).

Since heteromerization of PLN and SLN has been described and has been also suggested for other regulins (15), we investigated the hetero-oligomerization for PLN-ALN, PLN-ELN and ALN-ELN pairs in our system. Although heteromerization of these constructs was indicated by significantly increased FRET levels (Figure 4a), the values were dispersed. No or low FRET was observed in some of the cells and a markedly high FRET signal indicating tight protein-protein interaction was detected in other cells. This phenomenon was likely the result of the possible hetero-oligomerization combinations of two labeled proteins (Supplementary text). We also detected direct interaction between E protein and regulins, and the detected FRET signal was somewhat higher for E protein/PLN and E protein/ELN than for regulin heteromers (Figure 4a). The significance of these high values for E protein heteromers are underscored by the fact that only the E protein tagged at its N-terminal (luminal) end resulted in a stable fluorescent protein, thus eGFP and mCherry gave FRET signal in spite they localized at different sides of the ER membrane. It is worth mentioning that the thickness of the membrane bilayer is known to allow FRET signal detection (Supplementary text). We also observed cells with slightly increased FRET values for E protein and ALN coexpression, but this signal was not statistically significant.

**Figure 4.**
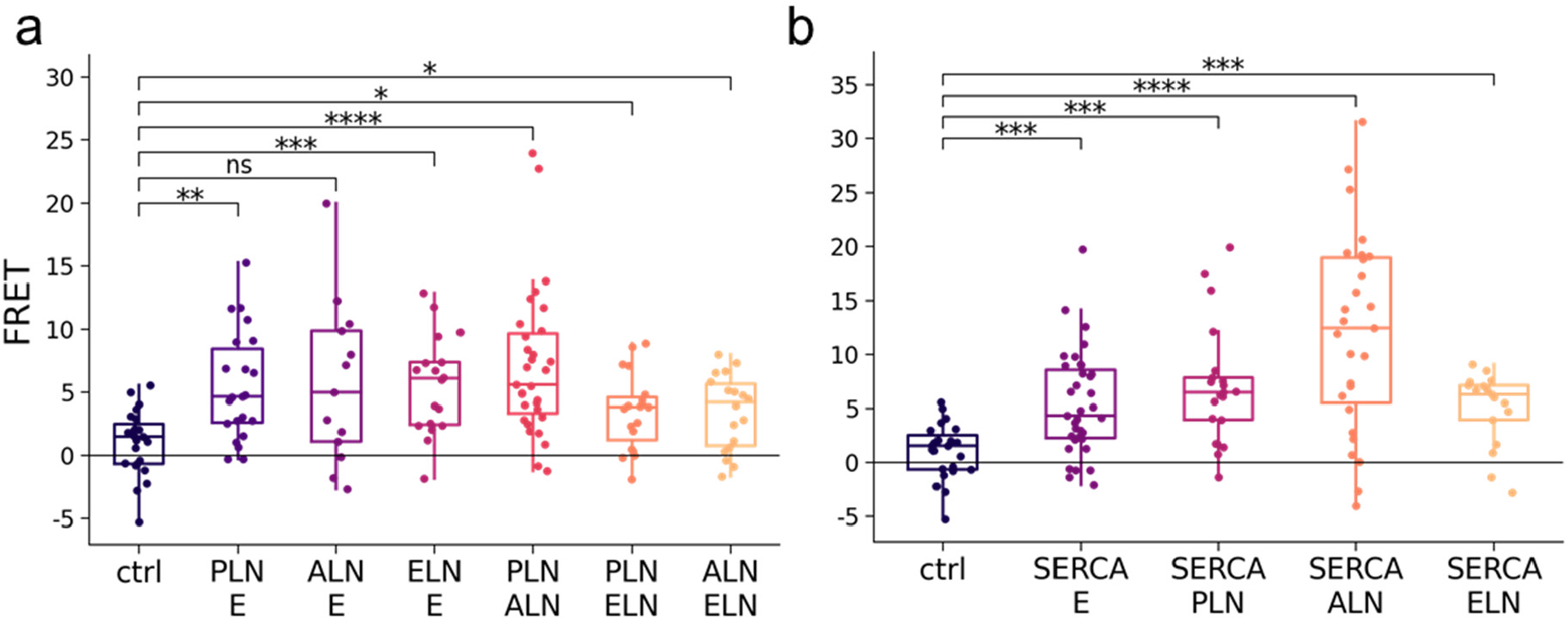
FRET efficiency for hetero-oligomerization among E protein, regulins (**a**), and SERCA (**b**). Individual data are shown as dots. Median and interquartile ranges are indicated with box plots. Data were analyzed by Kolmogorov-Smirnov test (* p<0.05, ** p<0.01, *** p<0.001, **** p<0.0001).

We also studied the E protein interaction with SERCA2b and found that the FRET values were significantly higher than those of the soluble eGFP and mCherry control (Figure 4b). When we compared the FRET values of E-protein/SERCA and regulins/SERCA interactions, we found that they were similar. The exception was the SERCA and ALN pair, which exhibited the highest FRET (Figure 4b). Importantly, the fluorescent proteins were also located on the opposite sides of the membrane bilayer in the case of the E protein/SERCA complex indicating strong interaction between these proteins.

### Structure and dynamics of E protein complexes

We aimed to characterize the structure and stability of complexes using the tools of 3D-bioinformatics and computational biology. Although we were able to use AlphaFold to predict the complexes of the plasma membrane Ca^2+^ ATPase (PMCA), which is a close relative of SERCA and its obligatory 1 TM partner proteins (PMCA/basigin and PMCA/neuroplastin) (27), AlphaFold was not able to build a rational structure for complexes including SERCA and regulins (Supplementary Figure S2). This failure may be a result of the transient and reversible nature of SERCA interactions, and a consequently low evolutionary information encoded on the protein-protein interaction interface, insufficient for successful AlphaFold predictions. Therefore, we used PIPER/ClusPro (28) to generate the structures of these complexes, and this resulted in docked E protein and regulins being located at the same site as observed in the SERCA/PLN experimental structure (Figure 5a-c).

**Figure 5.**
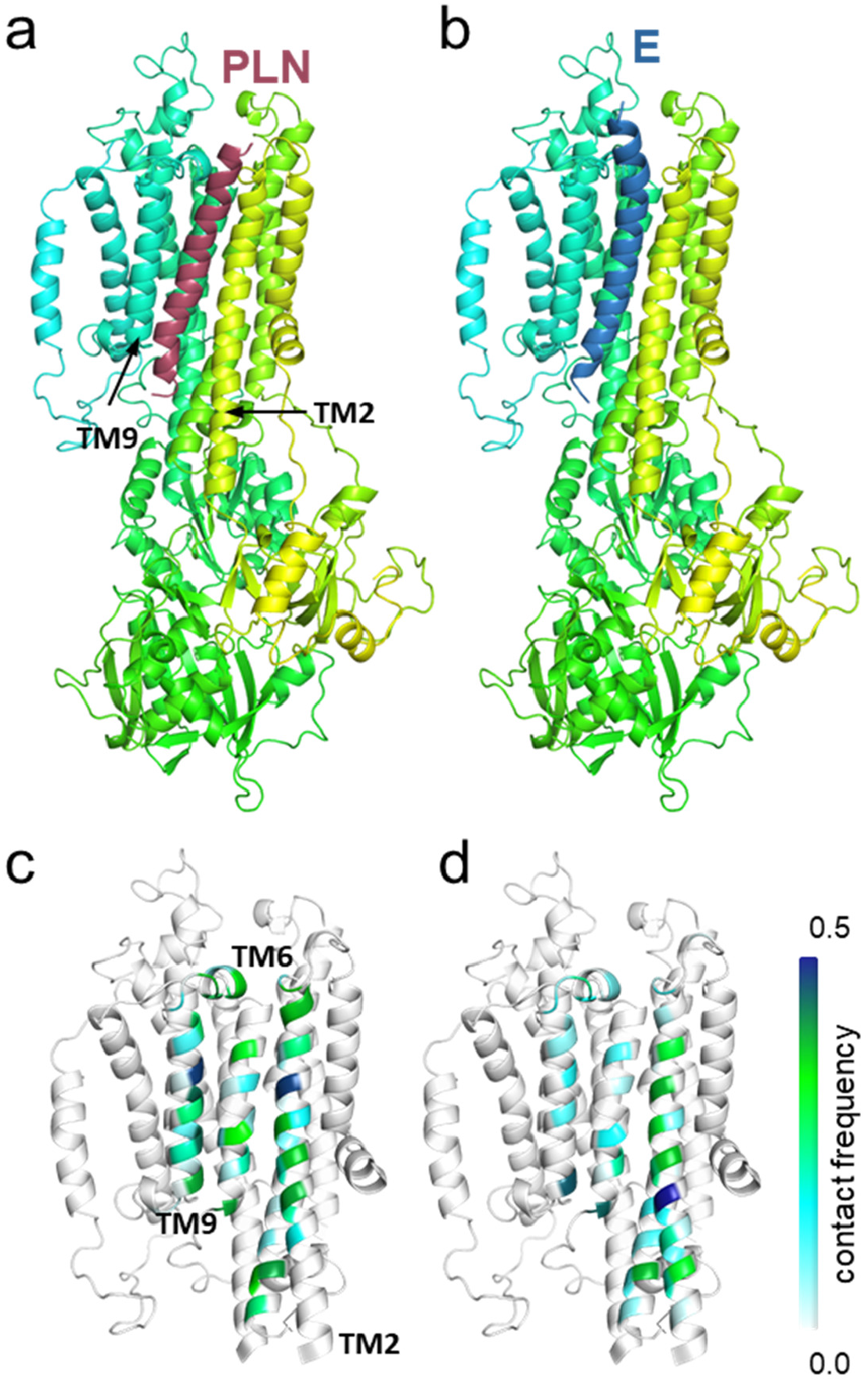
E protein binds to SERCA similarly to PLN. (**a**) PLN (burgundy) in the model of the human SERCA/PLN complex and (**b**) E protein (blue) from PIPER/ClusPro docking. SERCA is colored yellow-green-turquoise. The interaction frequency (white to blue) of (**c**) PLN and (**d**) E protein calculated from simulations are projected to SERCA TM helices.

In order to compare the stability of the E protein/SERCA complex to that of PLN/SERCA, we performed molecular dynamics (MD) simulations with these complexes (3 × 500 ns for each complex) embedded in a lipid bilayer. The E protein complex exhibited similar stability as the PLN/SERCA dimer and low fluctuations (Supplementary Figure S3). The PLN and E protein exhibited overlapping binding sites on SERCA. Both PLN and E protein interacted with SERCA TM2, TM6, and TM9, albeit in a slightly different manner (Figure 5e,f). E protein interactions were less frequent with most of the TM6 and TM9 residues and its most stable contacts were formed with residues at the intracellular bilayer boundary of these helices.

We also aimed to model the homo-and hetero-oligomers of regulins and E protein, and characterize their dynamics with MD simulations. However, we found that the E protein pentamer with a pore were unstable in simulations (Supplementary Figure S4) which was previously reported for regulins PLN and SLN (28).

### E protein causes unstable ER Ca^2+^ dynamics

To explore the functional significance of the interaction between E protein and SERCA2b, we investigated whether this interaction had an effect on calcium homeostasis. As a tracer for calcium signaling, we measured the ER Ca^2+^ level using the genetically encoded ER calcium indicator, ER-GCaMP6-150 (29). First, ATP was administered extracellularly to generate intracellular IP_3_ through purinergic receptors, leading to opening of the IP_3_ channels in the ER and to a decrease in the ER Ca^2+^ concentration. ATP was applied at two concentrations at 2 and 3 minutes of the experiments, resulting in three different responses in the case of HeLa cells with or without mCherry-tagged E protein (Figure 6a). We observed a switch-like response in several cells that caused a rapid and significant decrease in ER calcium levels which was frequently accompanied by Ca^2+^ oscillation. In control cells, this switch-like response occurred only in response to high ATP concentrations and was more prolonged than in cells expressing E protein. The second and third types of cellular response were the initiation of Ca^2+^ oscillation and a slow decrease in ER Ca^2+^ levels, respectively (Figure 6a).

**Figure 6.**
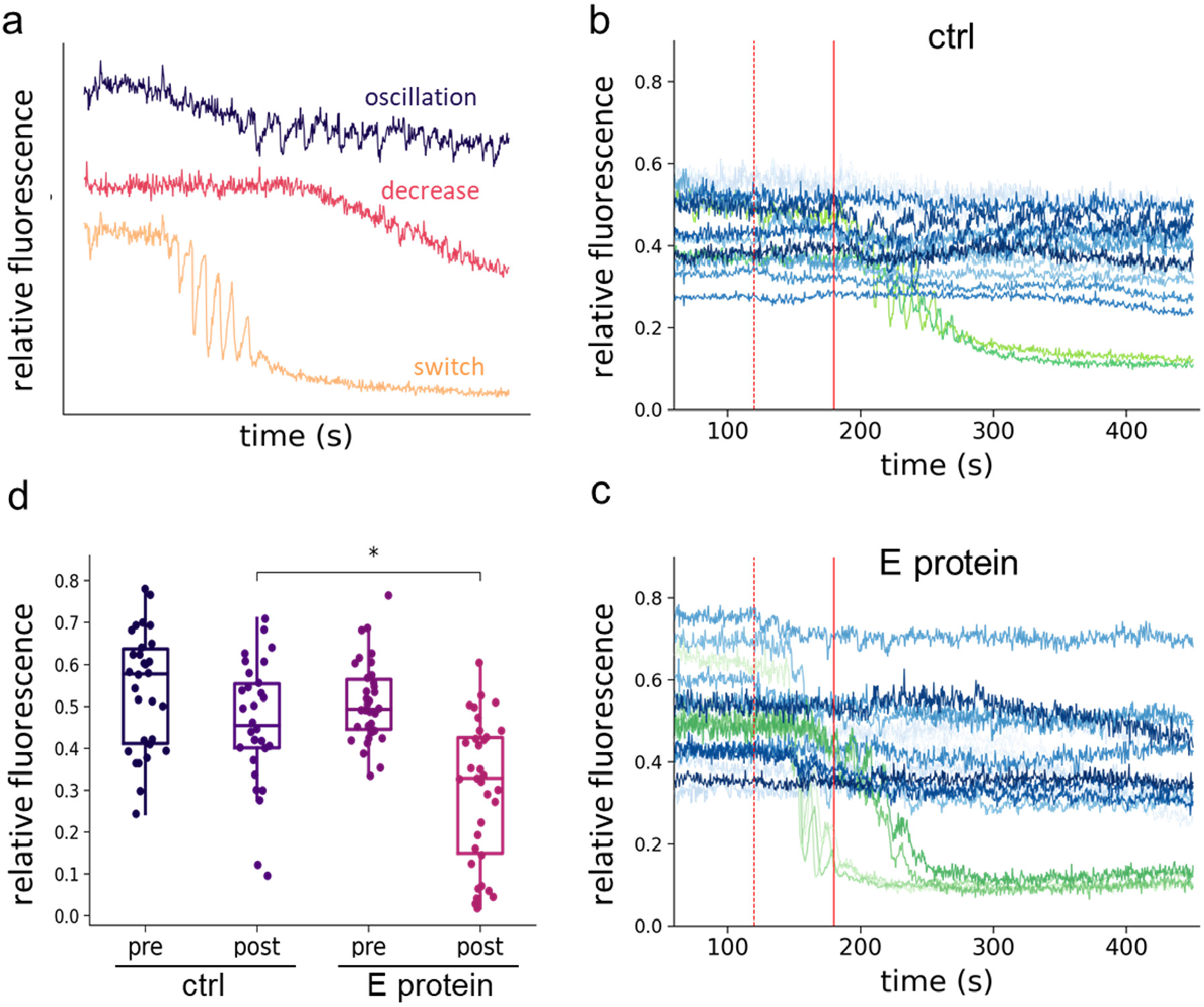
ER Ca^2+^ dynamics triggered by ATP. (**a**) Response types in the ER Ca^2+^ signal after addition of ATP. (**b, c**) Ca^2+^ level changes in cells expressing ER-GCaMP6-150 sensor alone (b) and with E protein (c) caused by ATP. Curves show the signal of individual cells. Red lines indicate ATP addition. (**d**) Ca^2+^ level before treatment (pre) and at 5 min after ATP addition (post). Data were analyzed by Wilcoxon–Mann–Whitney test (*** p<0.001).

We evaluated the change in ER Ca^2+^ levels by comparing the baseline levels before and 5 minutes after ATP administration. Following ATP administration, significantly lower Ca^2+^ levels were detected in the presence of E protein than in its absence (Figure 6b). In addition, strong switch-like responses corresponding to a sudden depletion of the ER store were observed more frequently in cells expressing E protein than in control cells. These results demonstrated that E protein made ER Ca^2+^ homeostasis metastable, sensitive to stimuli and depleting easily.

### E protein over-expression does not affect passive ER Ca^2+^ leakage but active reload

E protein is known to form homo-pentamers, providing a cation-selective pore (2). Since MD simulations suggested that the pore of the pentamer is not stable (Supplementary Figure S4), we investigated whether E protein was able to release Ca^2+^ ions from the ER into the cytoplasm. Therefore, we inhibited SERCA-mediated ER refill by thapsigargin (Tg) and monitored Ca^2+^ efflux from the ER, in the presence and absence of mCherry tagged E protein as indicated by the decrease in ER-GCaMP-150 fluorescence. The continuously decreasing ER Ca^2+^ levels measured in individual cells were fitted with exponential curves to obtain the leakage time constant (Figure 7). Since these time constants did not exhibit significant differences in the absence and presence of E protein, our results indicated that E protein did not affect the passive Ca^2+^ leakage from the ER suggesting that it did not conduct Ca^2+^ in our cellular system.

**Figure 7.**
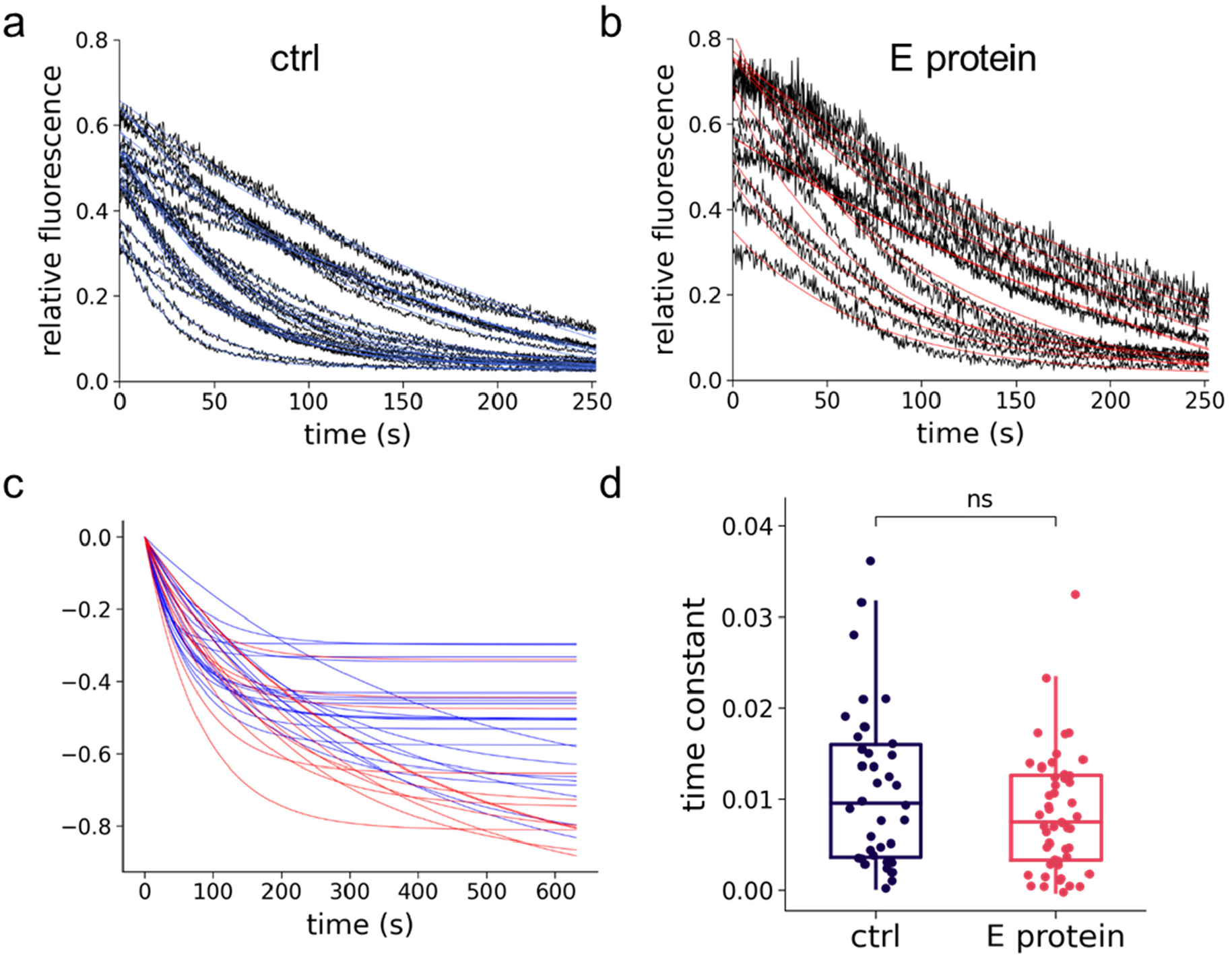
E protein does not alter the leakage from the ER. Ca^2+^ leakage from the ER after SERCA inhibition by thapsigargin, monitored with the ER-GCaMP6-150 Ca^2+^ sensor. The graphs show the change in [Ca^2+^]ER after Tg addition in HeLa cells expressing in the absence (control) **(a)** and in the presence of E protein **(b)**. Single exponential decay functions were fitted to the data points (blue and red curves) to determine the leakage time constant. **(c)** For comparison of leakage kinetics, the starting points of the fitted curves are shifted to zero. **(d)** Box plot of Ca^2+^ leakage time constants in control and E protein expressing cells (ns: not significant; Kolmogorov-Smirnov test).

Based on these observations, we examined with fluorescence microscopy whether E protein affected the ER Ca^2+^ reloading process. First, ER pools were emptied by ATP addition in a Ca^2+^-free medium. Then, the extracellular medium was replaced with a medium containing 2 mM Ca^2+^ to allow ER Ca^2+^ uptake. The kinetics of this uptake characterizes the pumping function of SERCA (30). Interestingly, the increase in the ER Ca^2+^ concentration was significantly slower and the time constant of this increase was significantly lower in cells expressing E protein than in the control cells (Figure 8). Since we found that E protein did not increase Ca^2+^ leakage, these results indicated an inhibition of SERCA-catalyzed ER Ca^2+^ refill by this viral protein. Importantly, this inhibition could be exerted only through the SERCA-system, as the SERCA pump was the only transport mechanism responsible for ER Ca^2+^ uptake. This was confirmed by the complete stop of ER reloading by using SERCA inhibitor Tg (Supplementary Figure S6). Since the plasma membrane was intact and the endoplasmic reticulum was emptied under identical conditions while monitoring the rate of release using the ER Ca2+ sensor, it is reasonable to assume that SERCA activity was monitored under identical Ca^2+^ gradients.

**Figure 8.**
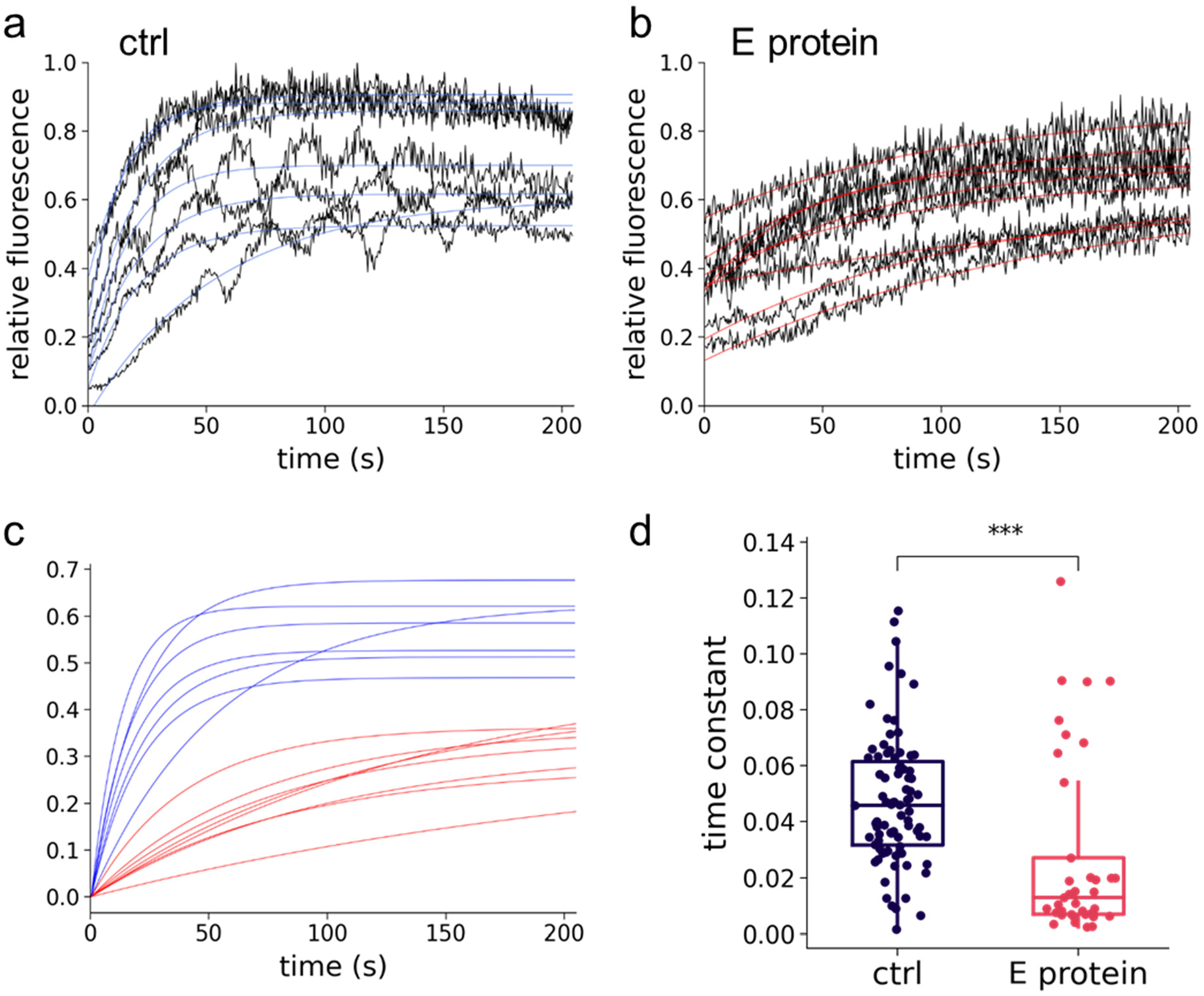
E protein alters the reload into the ER. Ca^2+^ reload into the ER after store depletion, achieved by ATP treatment in a calcium-free medium, was monitored after adding Ca^2+^ back to the medium in the absence (control) **(a)** and in the presence of E protein **(b)**. Single exponential decay functions were fitted to the data points (blue and red curves) to determine the reload time constant. **(c)** For comparison of reload kinetics, the starting points of the fitted curves are shifted to zero. (**d**) Box plot of Ca^2+^ reload time constants from control or E protein expressing individual cells. Median and interquartile range are indicated. Data were analyzed by Kolmogorov-Smirnov test (**** p<0.0001)

## DISCUSSION

Our results demonstrate that SARS-Cov-2 E protein interacts with members of the SERCA regulatory system, including SERCA2b as well as regulins based on FRET experiments. The relevance of SERCA/E protein interaction was confirmed by studying the Ca^2+^ dynamics of the ER. We found that in the presence of E protein, the ER Ca^2+^ signal was perturbed by a paracrine ATP trigger. This phenomenon was not due to a viroporin mediated Ca^2+^ channel function of the E protein, but rather an inhibition of SERCA by the E protein, reducing ER Ca^2+^ refill. Similar functional consequences of E protein on Ca^2+^ signaling in Figure 2 of Poggio *et al*. paper (31) can be observed, supporting our findings, however, those were not further explored by them.

We utilized AP-FRET and confirmed physical interaction between E protein, regulins and SERCA2b. Our results on regulin heterodimerization are in good agreement with a recent study on regulins with FRET-based experiments (15,25). Most importantly, our experiments demonstrated an interaction between E protein and SERCA confirming high-throughput assays, which identified this interaction (32,33). In addition, our MD simulations also supported the existence of the SERCA/E protein complex (Figure 5).

Interestingly, the highest FRET among SERCA-regulin pairs was measured for SERCA2b-ALN. Since the tissue and cell-type specific expressions of this pair exhibit a remarkable overlap, these proteins can be considered as a semi-obligate pair (23). To our knowledge, we investigated the SERCA2b isoform interactions with regulins using FRET for the first time. We highlight that the molecular-level structural matching of the different SERCA isoforms and regulins inferred from the FRET results is crucial, especially in systems such as the heart, where multiple SERCA isoforms (SERCA2a, SERCA2b) and multiple regulins (PLN, sarcolipin, ALN) are coexpressed (15).

E protein is known to form a pentameric structure providing an ion channel (34). Regulins are also known to form pentamers in the ER membrane creating a pool of oligomers, which is in dynamic equilibrium with the monomeric pool, directly involved in SERCA regulation. Therefore, our results on E protein/regulin heterodimerization strongly suggest that E protein can interfere with SERCA-regulation at multiple levels. Since the E protein pentamer collapsed in MD simulations (Supplementary Figure S4) and we did not observe ER Ca^2+^efflux through E protein in our experiments (Figure 7), we suppose that the E protein pore becomes stable only when other viral proteins are also expressed at a sufficient level. We find it unlikely that E protein concentration is low for channel forming in our heterologous expression system. Nevertheless, the delayed viroporin formation is possibly crucial for the virus to avoid early cell death before viral replication.

Importantly, the virally reprogrammed Ca^2+^ signaling can lead to different cell fates depending on the viral strain or the stage of viral replication. For example, Ca^2+^ depletion may delay apoptosis to gain time for replication (enteroviruses) (35) or it may promote apoptosis, facilitating virion release (hepatitis C) (36). Recently, the role of SERCA in viral infections has become a hot topic, related to the regulation of autophagy and inflammatory responses (20). Both of these processes were demonstrated to be influenced by SERCA regulation via Vacuole Membrane Protein 1 (VMP1) (20), which plays a key role in viral infections (20), also in SARS-Cov-2 infection (19). While the over-activation of the NLRP3 inflammasome is known to be responsible for severe COVID-19 cases (37) and viroporins are implicated in inflammasome activation (38), the molecular details of the involved processes are elusive. The VMP1, similarly to E protein, can interact with both SERCA and regulins (PLN and SLN), but these interactions activate SERCA, since VMP1 counteracts SERCA/regulin inhibitory interactions (39). It is also plausible that E protein interferes with SERCA regulation at the level of VMP1, as VMP1/E protein interaction has been observed in high throughput studies (40).

In summary, our results demonstrate that the SARS CoV-2 E protein modulates host cell Ca^2+^ homeostasis by affecting the SERCA-regulin system. The inhibitory effect was not exerted by Ca^2+^ conductance through E protein, but through a direct interaction between E protein and SERCA2b, possibly involving regulins. It is conceivable that important pathophysiological steps caused by viral infection, including autophagy and inflammatory responses, which are hallmarks of severe COVID-19 cases, may be rescued through the manipulation of SERCA activity. Therefore, it is of crucial importance to explore the infection associated chain of calcium signaling events in detail for enabling therapy.

## METHODS

### Constructs and chemicals

For mammalian expression of eGFP labelled proteins we used the pEGFP-C1 (Clontech) vector. For mCherry labelling we replaced eGFP to mCherry from pTK96_mCherry-MRLC2 (Addgene #46358) vector using AgeI and BsrGI sites. To construct the pEGFP-C1_SERCA vector carrying an N-terminally eGFP-labelled human SERCA2b, we amplified the SERCA2b cDNA from Addgene plasmid #75188 using SERCA specific forward 5’-GGGAGATCTATGGAGAACGCGCACACCAAGACGG-3’ and reverse 5’-GGGGTCGACTCAAGACCAGAACATATCGCTAAAGTTAG-3’ primers with overhanging BglII and SalI sites for pEGFP-C1 vector insertion. cDNAs of human regulins and SARS-CoV2 E protein with flanking BglII and BamHI restriction sites were custom synthesized (Integrated DNA Technologies; pIDTSMART-KAN vector; ALN: NM_001001701.3; ELN: NM_001162997.1; PLN: NM_002667.4, E protein: NC_045512.2). Vectors expressing N-terminally fluorescently labeled regulins and E protein were created by moving the cDNA of regulins into pEGFP-C1 and pmCherry-C vectors with BglII and BamHI.

ER-GCaMP6-150 cDNA was amplified from the Addgene plasmid #86918 using the ERGcAMPEcofw (5’-GGGGAATTCTCACAGCTCATCCTTG-3’) and ERGcAMPNotrev (5’-TTTGCGGCCGCATGGGACTGCTGTCT-3’) primers, digested with EcoRI and NotI, and inserted into the pSB-CMV-CAGPuro vector (gift of Tamas Orban). All constructs were verified by Sanger sequencing.

### Cell culture and transfection

HeLa cells were grown in Dulbecco’s modified Eagle’s medium (DMEM) supplemented with 10% Fetal Bovine Serum (FBS) and Penicillin/Streptomycin at 37°C in 5% CO_2_. The cells were seeded into an eight-well Nunc Lab-Tek II chambered cover glass (No:155409) at 4×10^4^/well density one day prior to transfection. Cells were transfected with the appropriate DNA constructs using FuGENE HD transfection reagent according to the manufacturer’s protocol. For FRET experiments, cells were fixed with 4% paraformaldehyde in PBS for 15min at 37°C.

### Fluorescent microscopy

For co-localization studies cells were washed with phosphate-buffered saline (PBS) and fixed with 4% paraformaldehyde in PBS for 15min at 37°C. Cells were permeabilized in pre-chilled (−20°C) methanol for 5°C. Samples were blocked for 1h at room temperature in PBS containing 2 mg/ml BSA, 0.1% Triton X-100 and 5% goat serum, then incubated for 1 h at room temperature with primary antibody (α-ERGIC; Sigma E1031) diluted in blocking buffer. After washing with PBS, cells were incubated for 1 h at room temperature with Alexa Fluor conjugated secondary antibody diluted 250 × in blocking buffer. After three washes, samples were stained with 5 μg/ml WGA Alexa Fluor 633 conjugate (10 min at room temperature). After repeated washes, samples were studied with a Nikon Eclipse Ti2.

### Acceptor-photobleaching (AP) FRET measurements

Using a Nikon Eclipse Ti2 confocal microscope with a 60x objective, images (PRE images) were taken in the channel corresponding to the donor (eGFP) and the acceptor (mCherry). The acceptor fluorophore was then quenched by high intensity light within a given area and new images were taken (POST images). To evaluate the images and to determine the FRET efficiency of the FRETcalc software was used to evaluate the images.

AP-FRET was performed with a Nikon Eclipse Ti2 confocal microscope with a 60x oil immersion objective. As donor, eGFP fluorophore was used, which was excited with the 488 nm line of a HeNe laser. The emission was collected between 505-550 nm. As acceptor, mCherry fluorophore was used, which was excited with the 561 nm laser line. The emission was collected above 580 nm. A region of interest (ROI) was selected within the ER compartment and 100% of 561 nm for 10 iterations was used for photobleaching the acceptor/mCherry to background levels. Pre-bleach and post-bleach images were acquired using identical imaging settings. FRET efficiency was calculated by the FRETcalc ImageJ plugin.

### Calcium signaling measurements

After 24 h of culture in 8-well chambered cover glass, cells were transfected with calcium indicator ER-GCaMP6-150 with or without mCherry-E protein. 24 h after transfection, medium was replaced by Hanks’s Buffered Salt Solution (HBSS, Thermo Fischer 88284) supplemented with 10 mM HEPES (pH 7.4) and 2 mM CaCl_2_.

Three types of experiments were performed. *a*, ATP stimulation: cells were stimulated with 240 μM and 400 μM ATP at 2 and 3 min, respectively. *b*, ER leakage: The extent of ER efflux was monitored under SERCA inhibition with 5 μM thapsigargin. *c*, The ER refill was measured using “Ca^2+^ re-addition” protocol with some modification (41). Intracellular Ca^2+^ stores were depleted by 400 μM ATP treatment in Ca^2+^-free HBSS supplemented with 10 mM HEPES (pH 7.4), 100 μM CaCl_2_ and 100 μM EGTA. After depletion, 2 mM Ca^2+^ was added and the progress of ER Ca^2+^ reload was monitored. Fluorescence changes of the ER-GCaMP-150 sensor were analyzed using the ImageJ Time Series Analyzer V3 plugin. The maximum and minimum values were obtained with 250 μM ionomycin and 2 mM EGTA, respectively, then the signals were normalized between 0 and 1.

### Structural models

Human SERCA/PLN homology model was built based on the rabbit complex (PDBID: 4kyt) and the human SERCA (PDBID: 7e7s) structures using Modeller (42) (Figure S3b and http://www.hegelab.org/resources.html). E protein was docked to the human SERCA model using PIPER/ClusPro (28). AlphaFold2 was run locally with full database mode as described in (43,44).

### Molecular dynamics simulations

The simulation systems were prepared using CHARMM-GUI (45,46). First, the structural models of SERCA/PLN and SERCA/E protein were oriented according to the Orientations of Proteins in Membranes database (47). Then N- and C-termini were patched with ACE (acetyl) and CT3 (N-Methylamide) groups, respectively. The membrane bilayer was symmetric containing 8:29:29:11:11:12 cholesterol:DMPC:POPC:DMPE:POPE:DMPI25 (Dimyristoyl-phosphatidylcholine; 1-Palmitoyl-2-oleoylphosphatidylcholine; Dimyristoyl-phosphatidylethanolamine; 1-Palmitoyl-2-oleoylphosphatidylethanolamine; Dimyristoyl-inositol-4,5-bisphosphate). KCl was used at a concentration of 150 mM. Grid information for PME (Particle-Mesh Ewald) electrostatics was generated automatically, and the number of particles, pressure of 1 bar, and temperature of 310 K were constant. GROMACS 2022 with the CHARMM36m force field was used to run molecular dynamics simulations (48,49). Each system was energy minimized using the steepest descent integrator, which stopped when the largest force in the system becomes less than 500 kJ/mol/nm. In order to increase sampling, several simulations (n=3 for each system) were forked using the energy minimized system, with different velocities. Equilibration was performed in six steps and production runs were run for 500 ns. The corresponding parameter files are also available for download. The trajectories were analyzed using MDAnalysis (50) and NumPy. Molecular visualization was performed using PyMOL (Schrödinger, LLC). Graphs were generated using Python and its matplotlib library (51).

## DATA AVAILABILITY

Some of our data are available at http://www.hegelab.org/resources.html or upon request.

## ACKNOWLEDGEMENTS

This research was funded by NRDIO/NKFIH grant numbers K127961, K137610, and TKP2021-EGA-23. We thank the computational resources made available on the GPU cluster Komondor (Governmental Information-Technology Development Agency, Hungary) and Erzsébet Suhajda for contributing to MD simulation analysis.

## ADDITIONAL INFORMATION

**Supplementary information** The online version contains supplementary material.

## REFERENCES

1. Polverino F, Kheradmand F. COVID-19, COPD, and AECOPD: Immunological, Epidemiological, and Clinical Aspects. Front Med (Lausanne). 2020;7:627278.

2. Schoeman D, Fielding BC. Coronavirus envelope protein: current knowledge. Virol J. 2019 May 27;16(1):69.

3. Weiss SR, Leibowitz JL. Coronavirus pathogenesis. Adv Virus Res. 2011;81:85–164.

4. Xia B, Shen X, He Y, Pan X, Liu FL, Wang Y, et al. SARS-CoV-2 envelope protein causes acute respiratory distress syndrome (ARDS)-like pathological damages and constitutes an antiviral target. Cell Res. 2021;31(8):847–60.

5. Venkatagopalan P, Daskalova SM, Lopez LA, Dolezal KA, Hogue BG. Coronavirus envelope (E) protein remains at the site of assembly. Virology. 2015 Apr;478:75–85.

6. Javorsky A, Humbert PO, Kvansakul M. Structural basis of coronavirus E protein interactions with human PALS1 PDZ domain. Commun Biol. 2021 Jun 11;4(1):724.

7. Jimenez-Guardeño JM, Nieto-Torres JL, DeDiego ML, Regla-Nava JA, Fernandez-Delgado R, Castaño-Rodriguez C, et al. The PDZ-binding motif of severe acute respiratory syndrome coronavirus envelope protein is a determinant of viral pathogenesis. PLoS Pathog. 2014;10(8):e1004320.

8. Surya W, Li Y, Torres J. Structural model of the SARS coronavirus E channel in LMPG micelles. Biochim Biophys Acta Biomembr. 2018 Jun;1860(6):1309–17.

9. Li Y, Surya W, Claudine S, Torres J. Structure of a conserved Golgi complex-targeting signal in coronavirus envelope proteins. J Biol Chem. 2014 May 2;289(18):12535–49.

10. Quaglia F, Salladini E, Carraro M, Minervini G, Tosatto SCE, Le Mercier P. SARS-CoV-2 variants preferentially emerge at intrinsically disordered protein sites helping immune evasion. FEBS J. 2022 Jul;289(14):4240–50.

11. Csizmadia G, Erdős G, Tordai H, Padányi R, Tosatto S, Dosztányi Z, et al. The MemMoRF database for recognizing disordered protein regions interacting with cellular membranes. Nucleic Acids Res. 2021;49(D1):D355–60.

12. Verdiá-Báguena C, Nieto-Torres JL, Alcaraz A, DeDiego ML, Torres J, Aguilella VM, et al. Coronavirus E protein forms ion channels with functionally and structurally-involved membrane lipids. Virology. 2012 Oct 25;432(2):485–94.

13. Wang WA, Carreras-Sureda A, Demaurex N. SARS-CoV-2 infection alkalinizes the ERGIC and lysosomes through the viroporin activity of the viral envelope protein. J Cell Sci. 2023 Mar 15;136(6):jcs260685.

14. Vangheluwe P, Sepúlveda MR, Missiaen L, Raeymaekers L, Wuytack F, Vanoevelen J. Intracellular Ca2+- and Mn2+-transport ATPases. Chem Rev. 2009 Oct;109(10):4733–59.

15. Singh DR, Dalton MP, Cho EE, Pribadi MP, Zak TJ, Šeflová J, et al. Newly Discovered Micropeptide Regulators of SERCA Form Oligomers but Bind to the Pump as Monomers. J Mol Biol. 2019;431(22):4429–43.

16. Peng J, Ran Y, Xie H, Deng L, Li C, Ling C. Sarco/Endoplasmic Reticulum Ca2+ Transporting ATPase (SERCA) Modulates Autophagic, Inflammatory, and Mitochondrial Responses during Influenza A Virus Infection in Human Lung Cells. J Virol. 2021 Apr 26;95(10):e00217-21, JVI.00217-21.

17. Kumar N, Khandelwal N, Kumar R, Chander Y, Rawat KD, Chaubey KK, et al. Inhibitor of Sarco/Endoplasmic Reticulum Calcium-ATPase Impairs Multiple Steps of Paramyxovirus Replication. Front Microbiol. 2019;10:209.

18. Ojha D, Basu R, Peterson KE. Therapeutic targeting of organelles for inhibition of Zika virus replication in neurons. Antiviral Res. 2023;209:105464.

19. Schneider WM, Luna JM, Hoffmann HH, Sánchez-Rivera FJ, Leal AA, Ashbrook AW, et al. Genome-Scale Identification of SARS-CoV-2 and Pan-coronavirus Host Factor Networks. Cell. 2021;184(1):120-132.e14.

20. Zack SR, Nikolaienko R, Cook B, Melki R, Zima AV, Campbell EM. Vacuole Membrane Protein 1 (VMP1) Restricts NLRP3 Inflammasome Activation by Modulating SERCA Activity and Autophagy. Res Sq. 2023;rs.3.rs-2508369.

21. Odermatt A, Taschner PE, Scherer SW, Beatty B, Khanna VK, Cornblath DR, et al. Characterization of the gene encoding human sarcolipin (SLN), a proteolipid associated with SERCA1: absence of structural mutations in five patients with Brody disease. Genomics. 1997;45(3):541–53.

22. Rathod N, Bak JJ, Primeau JO, Fisher ME, Espinoza-Fonseca LM, Lemieux MJ, et al. Nothing Regular about the Regulins: Distinct Functional Properties of SERCA Transmembrane Peptide Regulatory Subunits. Int J Mol Sci. 2021;22(16):8891.

23. Anderson DM, Makarewich CA, Anderson KM, Shelton JM, Bezprozvannaya S, Bassel-Duby R, et al. Widespread control of calcium signaling by a family of SERCA-inhibiting micropeptides. Sci Signal. 2016;9(457):ra119.

24. Cornea RL, Jones LR, Autry JM, Thomas DD. Mutation and phosphorylation change the oligomeric structure of phospholamban in lipid bilayers. Biochemistry. 1997 Mar 11;36(10):2960–7.

25. Phillips TA, Hauck GT, Pribadi MP, Cho EE, Cleary SR, Robia SL. Micropeptide heterooligomerization adds complexity to the calcium pump regulatory network. Biophys J. 2023;122(2):301–9.

26. Sudhaharan T, Ahmed S. Acceptor Photobleaching FRET (AP-FRET).

27. Naffa R, Hegedűs L, Hegedűs T, Tóth S, Papp B, Tordai A, et al. Plasma membrane Ca2+ pump isoform 4 function in cell migration and cancer metastasis. J Physiol. 2023 Mar 6;

28. Liu AY, Aguayo-Ortiz R, Guerrero-Serna G, Wang N, Blin MG, Goldstein DR, et al. Homologous cardiac calcium pump regulators phospholamban and sarcolipin adopt distinct oligomeric states in the membrane. Comput Struct Biotechnol J. 2022;20:380–4.

29. de Juan-Sanz J, Holt GT, Schreiter ER, de Juan F, Kim DS, Ryan TA. Axonal Endoplasmic Reticulum Ca2+ Content Controls Release Probability in CNS Nerve Terminals. Neuron. 2017 Feb 22;93(4):867-881.e6.

30. Chemaly ER, Troncone L, Lebeche D. SERCA control of cell death and survival. Cell Calcium. 2018;69:46–61.

31. Poggio E, Vallese F, Hartel AJW, Morgenstern TJ, Kanner SA, Rauh O, et al. Perturbation of the host cell Ca2+ homeostasis and ER-mitochondria contact sites by the SARS-CoV-2 structural proteins E and M. Cell Death Dis. 2023 Apr 29;14(4):297.

32. Samavarchi-Tehrani P, Abdouni H, Knight JDR, Astori A, Samson R, Lin ZY, et al. A SARS-CoV-2 – host proximity interactome [Internet]. bioRxiv; 2020 [cited 2023 Jun 9]. p. 2020.09.03.282103. Available from: https://www.biorxiv.org/content/10.1101/2020.09.03.282103v1

33. Nabeel-Shah S, Lee H, Ahmed N, Burke GL, Farhangmehr S, Ashraf K, et al. SARS-CoV-2 nucleocapsid protein binds host mRNAs and attenuates stress granules to impair host stress response. iScience. 2022 Jan 21;25(1):103562.

34. Cao Y, Yang R, Lee I, Zhang W, Sun J, Wang W, et al. Characterization of the SARS-CoV-2 E Protein: Sequence, Structure, Viroporin, and Inhibitors. Protein Sci. 2021 Jun;30(6):1114–30.

35. van Kuppeveld FJM, de Jong AS, Melchers WJG, Willems PHGM. Enterovirus protein 2B po(u)res out the calcium: a viral strategy to survive? Trends Microbiol. 2005 Feb;13(2):41–4.

36. Kominek J, Doering DT, Opulente DA, Shen XX, Zhou X, DeVirgilio J, et al. Eukaryotic Acquisition of a Bacterial Operon. Cell. 2019 Mar 7;176(6):1356-1366.e10.

37. Kaivola J, Nyman TA, Matikainen S. Inflammasomes and SARS-CoV-2 Infection. Viruses. 2021 Dec;13(12):2513.

38. Guarnieri JW, Angelin A, Murdock DG, Schaefer P, Portluri P, Lie T, et al. SARS-COV-2 viroporins activate the NLRP3-inflammasome by the mitochondrial permeability transition pore. Frontiers in Immunology [Internet]. 2023 [cited 2023 Jun 9];14. Available from: https://www.frontiersin.org/articles/10.3389/fimmu.2023.1064293

39. Zhao YG, Chen Y, Miao G, Zhao H, Qu W, Li D, et al. The ER-Localized Transmembrane Protein EPG-3/VMP1 Regulates SERCA Activity to Control ER-Isolation Membrane Contacts for Autophagosome Formation. Mol Cell. 2017 Sep 21;67(6):974-989.e6.

40. St-Germain JR, Astori A, Samavarchi-Tehrani P, Abdouni H, Macwan V, Kim DK, et al. A SARS-CoV-2 BioID-based virus-host membrane protein interactome and virus peptide compendium: new proteomics resources for COVID-19 research [Internet]. bioRxiv; 2020 [cited 2023 Jun 13]. p. 2020.08.28.269175. Available from: https://www.biorxiv.org/content/10.1101/2020.08.28.269175v1

41. Pászty K, Caride AJ, Bajzer Ž, Offord CP, Padányi R, Hegedűs L, et al. Plasma membrane Ca^2+^-ATPases can shape the pattern of Ca^2+^ transients induced by store-operated Ca^2+^ entry. Sci Signal. 2015 Feb 17;8(364):ra19.

42. Fiser A, Sali A. Modeller: generation and refinement of homology-based protein structure models. Methods Enzymol. 2003;374:461–91.

43. Tordai H, Suhajda E, Sillitoe I, Nair S, Varadi M, Hegedus T. Comprehensive Collection and Prediction of ABC Transmembrane Protein Structures in the AI Era of Structural Biology. Int J Mol Sci. 2022;23(16):8877.

44. Hegedűs T, Geisler M, Lukács GL, Farkas B. Ins and outs of AlphaFold2 transmembrane protein structure predictions. Cell Mol Life Sci. 2022;79(1):73.

45. Jo S, Kim T, Iyer VG, Im W. CHARMM-GUI: a web-based graphical user interface for CHARMM. J Comput Chem. 2008;29(11):1859–65.

46. Wu EL, Cheng X, Jo S, Rui H, Song KC, Dávila-Contreras EM, et al. CHARMM-GUI Membrane Builder toward realistic biological membrane simulations. J Comput Chem. 2014 Oct 15;35(27):1997–2004.

47. Lomize MA, Lomize AL, Pogozheva ID, Mosberg HI. OPM: orientations of proteins in membranes database. Bioinformatics. 2006 Mar 1;22(5):623–5.

48. Huang J, Rauscher S, Nawrocki G, Ran T, Feig M, de Groot BL, et al. CHARMM36m: an improved force field for folded and intrinsically disordered proteins. Nat Methods. 2017 Jan;14(1):71–3.

49. Van Der Spoel D, Lindahl E, Hess B, Groenhof G, Mark AE, Berendsen HJC. GROMACS: fast, flexible, and free. J Comput Chem. 2005;26(16):1701–18.

50. Michaud-Agrawal N, Denning EJ, Woolf TB, Beckstein O. MDAnalysis: a toolkit for the analysis of molecular dynamics simulations. J Comput Chem. 2011 Jul 30;32(10):2319–27.

51. Hunter JD. Matplotlib: A 2D Graphics Environment. Computing in Science & Engineering. 2007;9(3):90–5.

