## Supplementary Material for "SARS-CoV-2 Envelope protein alters calcium signaling via SERCA interactions"

### Supplementary text

Heterogeneity of FRET values. In general, we recorded FRET values with a large variability. There were some cells in which no FRET was detected at all, and others in which particularly high FRET was observed. One reason for this in the case of regulins and E protein is their different subcellular localization (e.g. trafficking from ER to Golgi), so their intracellular distribution can be different from cell to cell. Interestingly, we observed quite different trafficking patterns for different regulins. For example, ALN seemed to reach the Golgi localization shortly after transfection. On the other hand, both regulins and E protein can homo-oligomerize with themselves or form hetero-oligomers with the others and the different oligomerization modes can compete with each other. Moreover, the amount of proteins in the cells and the proportion of proteins with the different fluorescence tags may also vary during co-expression.

FRET through a membrane bilayer. The physical interaction between E protein, regulins and SERCA were detected using AP-FRET with N-terminally labeled fluorescent constructs. The N-termini of both regulins and SERCA are cytosolic, while the E protein has opposite orientation with a luminal N-terminus (1). Since the C-terminally tagged E protein was not stable, we used N-terminal labels and detected energy transfer between the two fluorophores separated by a lipid bilayer. A similar setup was described previously for both plasma membrane (2,3) and endoplasmic reticulum membranes confirming the feasibility of our assay (4). While the ER membrane thickness can vary between 2.9-5.5 nm depending on the lipid composition (5), the fluorescence energy transferred from a donor fluorophore to an acceptor fluorophore is typically over distances between 1 nm and 10 nm (6). In addition, the E protein N-terminus is short, ending near the membrane/cytosol interface, thus it restrains the fluorescent protein close to the membrane. These observations support the validity of our FRET results.

**Immunostaining and fluorescent microscopy**

For co-localization studies, the endogenous ERGIC-53 protein was stained by the α-ERGIC-53 antibody (Sigma, E1031; ERGIC marker) and the ER $Ca^{2+}$ sensor, ER-GCaMP6-150 was stained by α-GFP (Invitrogen, MA5-15256; ER marker). One day after transfection, cells were washed with phosphate-buffered saline (PBS) and fixed with 4% paraformaldehyde in PBS for 15 min at 37ºC. Cells were permeabilized in pre-chilled (-20ºC) methanol for 5ºC. Samples were blocked for 1h at room temperature in PBS containing 2 mg/ml BSA, 0.1% Triton X-100 and 5% goat serum, then incubated for 1 h at room temperature with primary antibodies diluted in blocking buffer. After washing with PBS, cells were incubated for 1 h at room temperature with Cy3 conjugated secondary antibody diluted 250 × in blocking buffer. After repeated washes, samples were studied with a Nikon Eclipse Ti2 using 60x oil immersion objective. Green and red fluorescence was acquired at 505-550 nm and >580 nm using excitations at 488 and 561 nm laser lines, respectively.

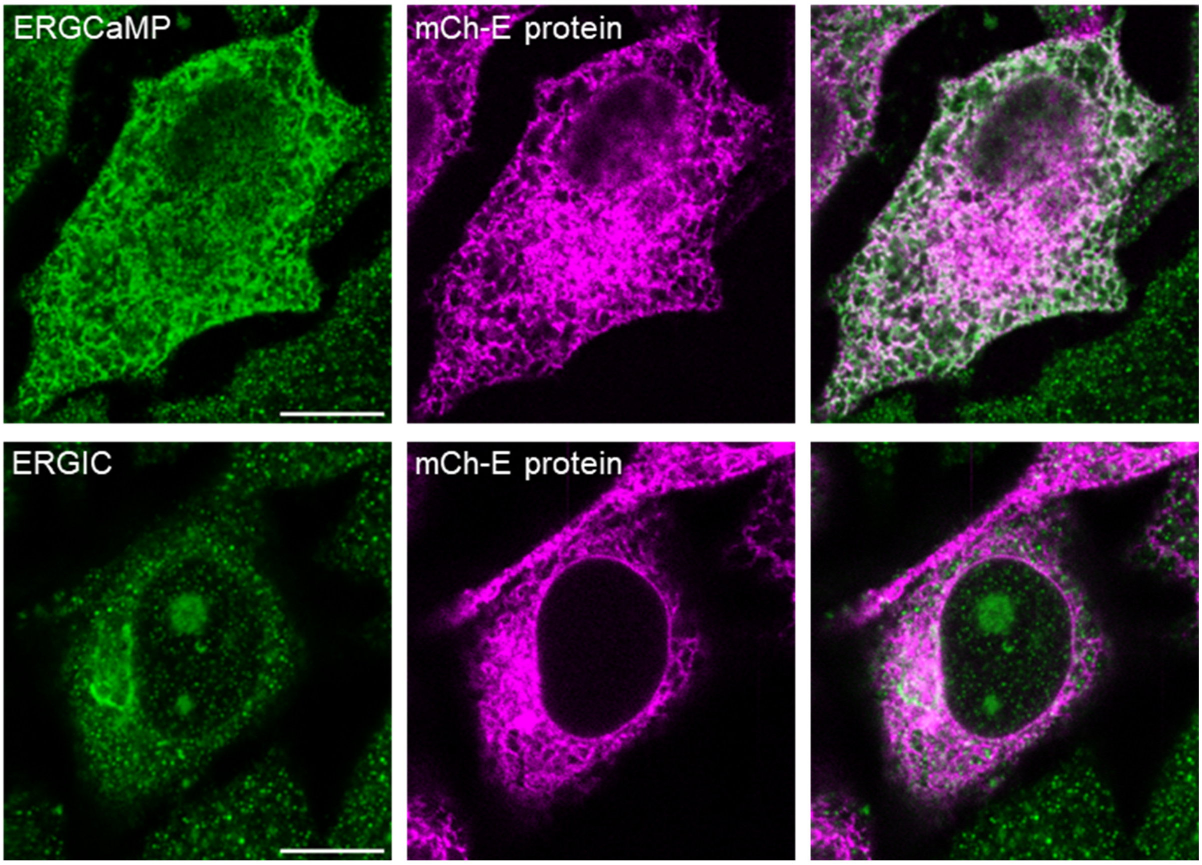


**Supplementary Figure S1:** mCherry-E protein colocalizes with an ER marker (ER-GCaMP6-150) and an ERGIC marker (α-ERGIC-53) in HeLa cells. In the first column ER and ERGIC markers are shown in green. Second column displays the distribution of mCherry-E protein. Third column exhibits the two channels together. Colocalized proteins are indicated by white color.

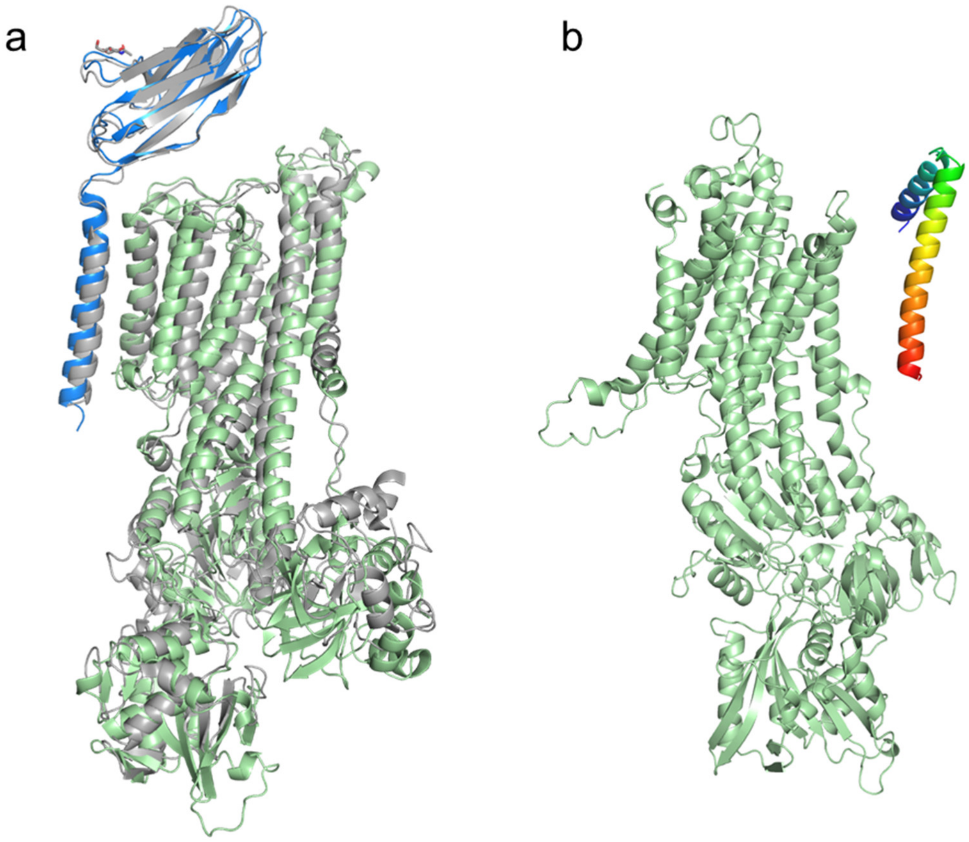


**Supplementary Figure S2:** (**a**) AlphaFold correctly predicted the PMCA/neuroplastin complex (green and blue) when compared to its experimental structure (gray, PDBID: 6a69) (7,8). (**b**) SERCA and PLN do not interact in the AlphaFold-predicted structure. The failure was likely caused by the low level of evolutionary information associated with this transient protein-protein interaction.

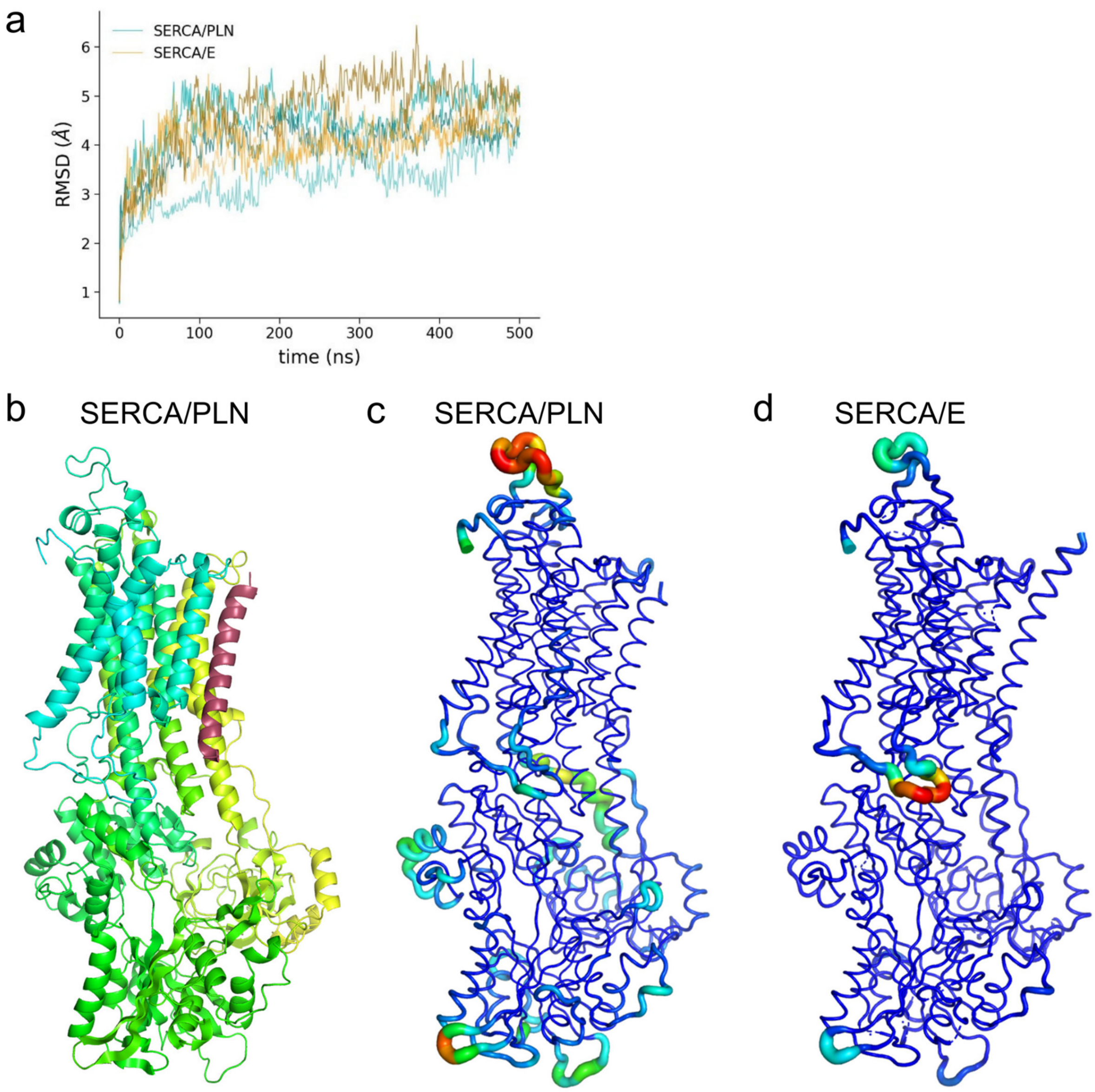


**Supplementary Figure S3:** (**a**) RMSD (**b**) Human SERCA/PLN homology model based on the rabbit complex (PDBID: 4kyt) and the human SERCA (PDBID: 7e7s) structures. SERCA is colored yellow-green-cyan and PLN is labeled by raspberry. (**c, d**) Structures in rainbow colors show the root mean square fluctuations calculated using GROMACS Tools and plotted using PyMOL. Warmer color and thicker representation indicate higher fluctuations.

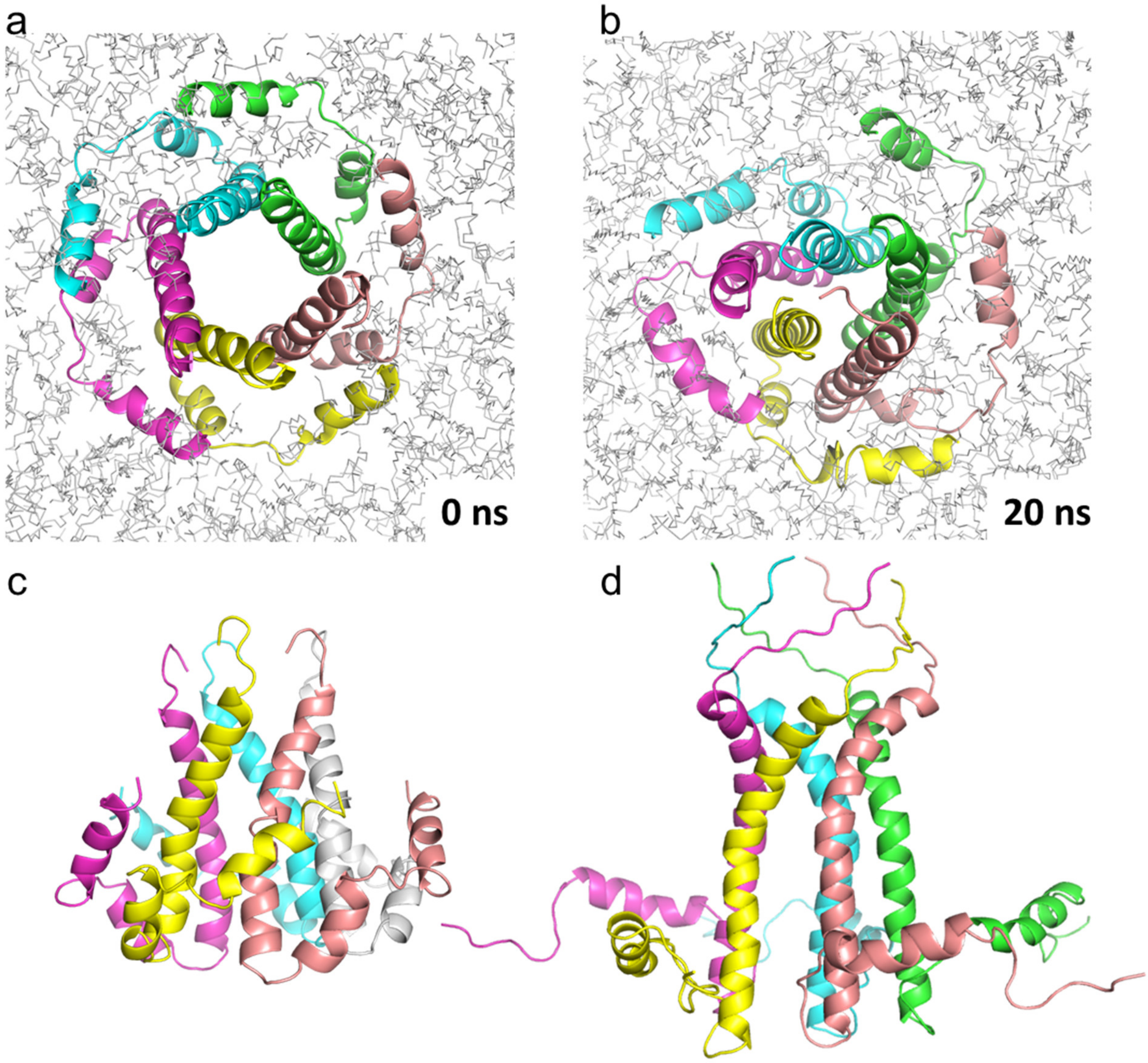


**Supplementary Figure S4:** (**a, b**) E protein channel collapsed fast in MD simulations with structure from SARS-CoV-1 (PDBID: 5x29). (**c**) The NMR structure of this pentamer exhibited C-terminal regions immersed in the lipid bilayer. To exclude this conformation as a source of the collapse, we generated a simulation system only with TM regions (not shown), but it exhibited similar instability. (**d**) A pentamer was modeled based on a PLN homopentamer (PDBID: 2m3b), since the C-terminal helices in this structure were aligned in parallel with the membrane bilayer. However, the central channel was also not stable in this case and the luminal parts became highly bent.

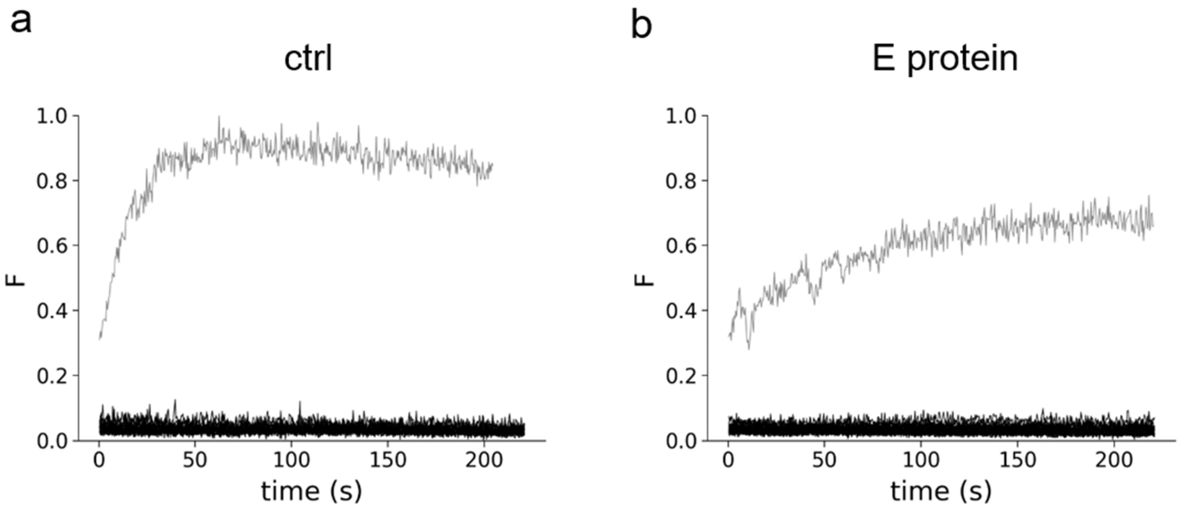


**Supplementary Figure S5:** Inhibition of SERCA activity by thapsigargin completely abolished the ER reloading. Experiments of $Ca^{2+}$ reload into the ER were repeated with SERCA inhibitor thapsigargin (5 μM) in the absence (**a**) or in the presence of E protein (**b**). Store depletion was achieved by ATP treatment in a calcium-free medium, and the ER $Ca^{2+}$ concentration was monitored after adding $Ca^{2+}$ back to the medium (black curves, $n_{cntr}$=17, $n_{Eprotein}$=24). An example curve for $Ca^{2+}$ reload in the presence of active SERCA without inhibition is shown in gray from Figure 8.

**Supplementary references**


1. Nieto-Torres JL, DeDiego ML, Álvarez E, Jiménez-Guardeño JM, Regla-Nava JA, Llorente M, et al. Subcellular location and topology of severe acute respiratory syndrome coronavirus envelope protein. Virology. 2011 Jul 5;415(2):69–82.

2. Sappakhaw K, Jantarug K, Slavoff SA, Israsena N, Uttamapinant C. A Genetic Code Expansion-Derived Molecular Beacon for the Detection of Intracellular Amyloid-β Peptide Generation. Angewandte Chemie International Edition. 2021;60(8):3934–9.

3. Haga Y, Ishii K, Hibino K, Sako Y, Ito Y, Taniguchi N, et al. Visualizing specific protein glycoforms by transmembrane fluorescence resonance energy transfer. Nat Commun. 2012 Jun 19;3(1):907.

4. Fernández-Dueñas V, Burgueño J, Ciruela F. Exploring Drug-Receptor Interaction Kinetics: Lessons from a Sigma-1 Receptor Transmembrane Biosensor. Frontiers in Pharmacology [Internet]. 2017 [cited 2023 Jun 9];8. Available from: https://www.frontiersin.org/articles/10.3389/fphar.2017.00004

5. Prasad R, Sliwa-Gonzalez A, Barral Y. Mapping bilayer thickness in the ER membrane. Sci Adv. 2020;6(46):eaba5130.

6. Algar WR, Hildebrandt N, Vogel SS, Medintz IL. FRET as a biomolecular research tool - understanding its potential while avoiding pitfalls. Nat Methods. 2019 Sep;16(9):815–29.

7. Gong D, Chi X, Ren K, Huang G, Zhou G, Yan N, et al. Structure of the human plasma membrane Ca2+-ATPase 1 in complex with its obligatory subunit neuroplastin. Nat Commun. 2018 Sep 6;9(1):3623.

8. Naffa R, Hegedűs L, Hegedűs T, Tóth S, Papp B, Tordai A, et al. Plasma membrane Ca2+ pump isoform 4 function in cell migration and cancer metastasis. J Physiol. 2023 Mar 6;
